# Tubulin autoregulation tunes microtubule dynamics to support multicellular architecture and viability

**DOI:** 10.1101/2025.07.28.667019

**Authors:** Ana C. Almeida, Chahrazed Lacheheub, Ivana Gasic

**Affiliations:** Department of Molecular and Cellular Biology, University of Geneva, Geneva, Switzerland

## Abstract

Alpha- and beta-tubulin heterodimers dynamically assemble into microtubules, key cytoskeletal elements involved in intracellular trafficking, cell adhesion and division. The availability of free tubulins regulates the synthesis of new subunits. In response to excessive soluble αβ-tubulins, tetratricopeptide protein 5 (TTC5) selectively recognizes nascent tubulins at the ribosome, recruiting downstream effectors that degrade their encoding messenger RNAs, in a process known as tubulin autoregulation. Despite its well-characterized molecular framework, the biological relevance of this regulatory pathway remains unknown. Here, using human 3D cellular models, advanced optics, and genetic perturbation of tubulin biosynthesis, we reveal that loss of TTC5-dependent tubulin autoregulation elevates soluble tubulin levels, inducing microtubule hyperstability, and disrupting cytoskeletal organization. These defects impair the localization of adhesion molecules at cell-cell junctions and extracellular matrix interfaces, compromising tissue architecture and reducing overall cell viability. Our findings establish tubulin autoregulation as a critical mechanism that tunes microtubule dynamics to sustain cellular integrity and tissue homeostasis.

## Introduction

Microtubules, composed of α- and β-tubulin heterodimers (henceforth αβ-tubulins), are key cytoskeletal elements that provide structural support, facilitate intracellular transport, and ensure accurate cell divison^1–3^. These functions critically depend on the precise yet evolving control of αβ-tubulin levels and their spatial distribution. The best characterized microtubule regulatory mechanism is their intrinsic dynamic instability, manifested by stochastic exchanges of polymerized and soluble αβ-tubulin subunits^4^. This property, regulated by a plethora of microtubule-associated proteins, allows continuous remodeling of microtubule cytoskeleton^5–7^. Much less understood is how cells define and maintain the correct supply of αβ-tubulin proteins needed to build a microtubule network tailored to cellular demands.

Evidence from *in vitro* work suggests that microtubule nucleation, polymerization, and dynamics scale with the concentration of αβ-tubulins^8–11^. Whether this also holds true in cells remains unknown. The existence of a conserved and sophisticated feedback mechanism that controls tubulin biosynthesis suggests that maintaining precise αβ-tubulin abundance is biologically relevant.

When in surplus, soluble αβ-tubulins trigger a pathway named tubulin autoregulation, which selectively destabilizes tubulin-encoding mRNAs^12,13^. This process is mediated by TTC5 that specifically recognizes nascent α- and β-tubulins at the ribosomal exit tunnel via their conserved amino-terminal tetrapeptides^14–17^. Upon recognition, TTC5 recruits the adaptor protein SCAPER (S-Phase Cyclin A Associated Protein in the ER) and effector CCR4-NOT complex (Carbon Catabolite Repression-Negative On TATA-less) to tubulin-translating ribosomes, initiating mRNA deadenylation and decay^18,19^. Soluble αβ-tubulins directly input into pathway activity via reversible sequestration of TTC5: under steady-state conditions, αβ- tubulins bind and maintain TTC5 in an inactive state. When αβ-tubulin levels rise, TTC5 is released and activates tubulin autoregulation machinery^20^.

Although tubulin autoregulation was discovered over four decades ago, its core molecular mediators have only recently been identified^18–20^. The biological function of tubulin autoregulation, however, remains largely unexplored. Experimental activation of this pathway has historically relied on artificially elevating soluble αβ-tubulins using microtubule-depolymerizing agents such as colchicine^12,13^, or direct tubulin injection^21^, thereby obscuring its physiological role.

Altered expression and mutations in core mediators of tubulin autoregulation, TTC5 and SCAPER, have been associated with diverse cancers (*COSMICv101*^22^, *GRCh38*), ciliopathies and neurodevelopmental disorders, such as tubulinopathies^23–26^, hinting that tubulin autoregulation may play a role in maintaining genome stability and cellular homeostasis.

Tubulin mRNA levels fluctuate in response to various physiological and pathological stimuli, including nutrient deprivation or oncogenic transformation, suggesting that tubulin gene expression dynamically adapts to cellular demands^27^. Surprisingly, studies in human cultured cells lacking tubulin autoregulation reported no detectable changes in tubulin levels^18^, despite clear chromosome segregation defects during cell division^18–20^. These findings were conducted in 2D bulk cell cultures, where subtle, cell-cycle-specific, or context-dependent effects may be masked. Indeed, tubulin expression is influenced by culture conditions, including spatial organization (2D versus 3D)^27^ and interactions with extracellular matrix (ECM)^28^, emphasizing the need to assess tubulin biosynthesis control in more complex tissue-like models.

While conventional 2D cell cultures have been instrumental in advancing our understanding of microtubule biology, they offer limited spatial and mechanical cues, which can influence cell adhesion, shape, and protein dynamics^29,30^. To better recapitulate cell-cell interactions and microenvironmental gradients present *in vivo*, we employ 3D spheroid models. Spheroids provide a reproducible and tractable system to study fundamental biological problems in a multicellular context^31,32^. Their simplicity, scalability, and amenability to genetic manipulation make them well suited for functional dissection of tubulin autoregulation and its role in maintaining tissue-like architecture and integrity.

## Results

### Tubulin autoregulation maintains steady-state tubulin levels and microtubule dynamics

To explore tubulin autoregulation in a multicellular model system, we leveraged HeLa cells that self-assemble into spheroids during proliferation, reliant on extensive cell-cell and cell-ECM interactions^33^. Spheroids typically organize into a stratified structure: a proliferative outer rim, a quiescent middle layer, and a necrotic core caused by oxygen and nutrient limitations^34,35^ (Fig. 1A). We captured distinct HeLa spheroid development phases using brightfield imaging: spheroid aggregation and compaction (days 0-2), and growth (days 5-10)^36,37^ (Fig. 1B).

**Fig. 1.**
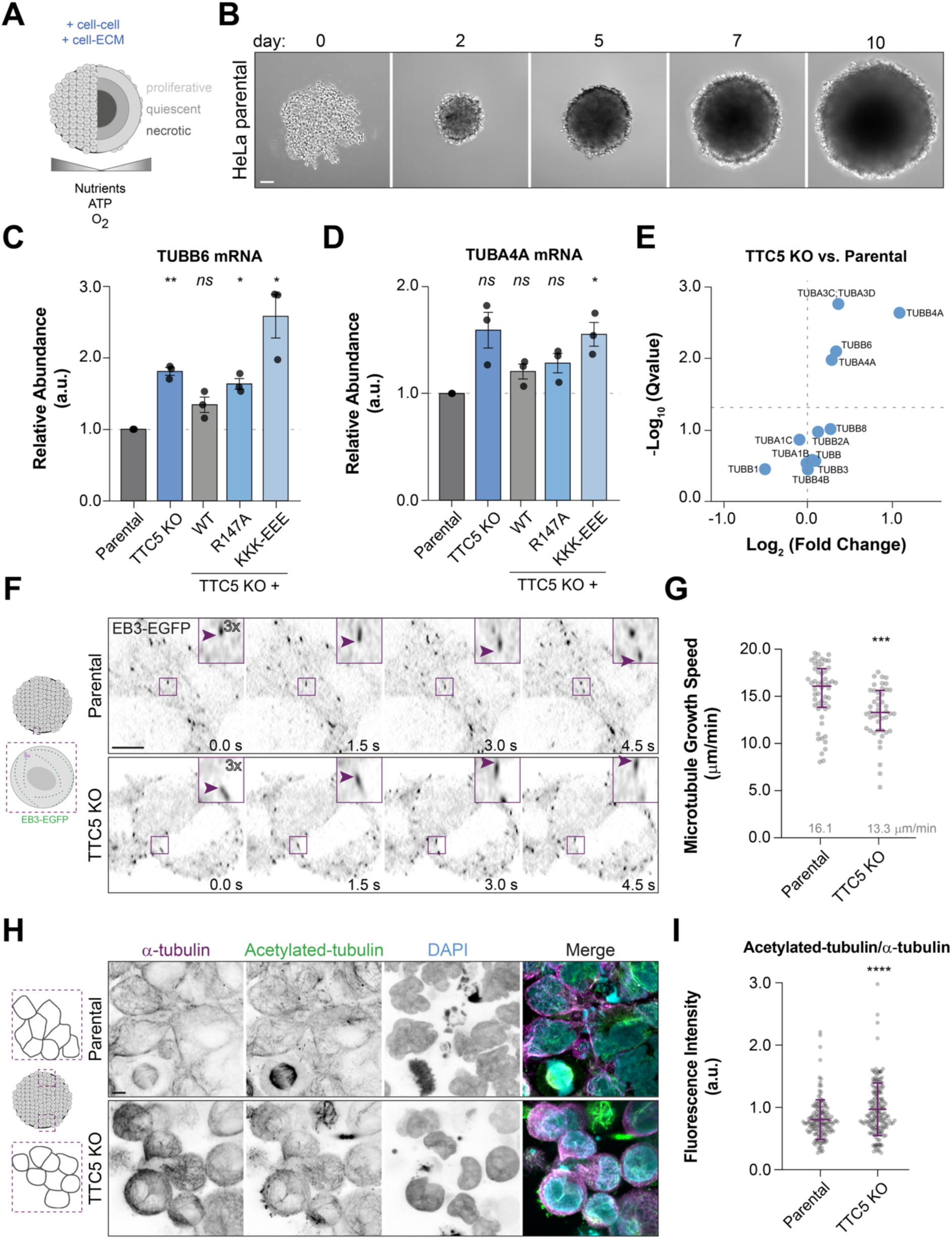
Tubulin autoregulation maintains steady-state tubulin levels and microtubule dynamics. (A) Schematic representation of the spheroid model. (B) Representative brightfield images of HeLa parental spheroids cultured for 10 days. Scale bar, 100 μm. (C) Relative abundance of TUBB6 and (D) TUBA4A mRNA in HeLa parental, TTC5 knockout, and the indicated Flag-TTC5 cell lines in 10-days old spheroids, normalized to housekeeping transcripts and the parental levels. Data show mean ± SD from three independent experiments. *p* values indicate unpaired, two-tailed Student’s t-test for each of the indicated cell lines against parental cell line. (E) Volcano-plot of the relative abundance of α- and β-tubulin isotypes in TTC5 KO versus parental spheroids, measured by quantitative mass spectrometry. Horizontal dash line indicates *q*-value of 0.05. (F) Time-lapse imaging of EB3-EGFP in HeLa parental and TTC5 KO cells, showing microtubule plus-end tracking (1.5 second interval). Magenta arrowhead follows a representative microtubule growth event over time. Scale bar, 5 μm. (G) Microtubule growth speed in interphase parental and TTC5 KO cells. Data show median ± interquartile range from three independent experiments. *p* value indicates unpaired, unpaired, Mann-Whitney test comparing ranks between TTC5 KO to the parental cell line. (H) Immunofluorescence of 10-day HeLa parental and TTC5 KO spheroids stained for total α-tubulin (magenta), acetylated α-tubulin (green) and DNA (DAPI, cyan). Scale bar, 5 μm. (I) Quantification of acetylated α-tubulin/total α-tubulin intensity at the cell cortex (defined by actin localization, see Extended Data Fig. 1H). Data are median ± interquartile range from three independent experiments. *p* value from unpaired, Mann-Whitney test comparing ranks of TTC5 KO with parental cells. *\*\*\*\*p*<0.0001, *\*\*\*p*<0.001, *\*\*p*<0.01, *\*p<*0.05, *ns* - not significant.

We first confirmed that tubulin autoregulation responds to its known activator in spheroids by quantifying mRNA levels upon 6 hours colchicine treatment. TUBA and TUBB transcripts decreased in parental and TTC5 knockout spheroids that re-expressed wild type TTC5 (TTC5^WT^), but not in TTC5 knockout (KO) or spheroids expressing TTC5 mutants that abolish its function in tubulin mRNA decay: the nascent tubulin peptide- and ribosome-binding mutant (TTC5^R147A^ and TTC5^KKK-EEE^, henceforth autoregulation-incompetent mutants) (Extended Data Fig. 1A). These results mirror previous reports in 2D cell cultures^18^.

Strikingly, even untreated 10-days old TTC5 KO and autoregulation-incompetent mutant spheroids showed a reproducible increase in tubulin mRNA and protein levels, as measured by RT-qPCR (Fig. 1C-D, Extended Data Fig. 1B-C) and immunoblotting (Extended Data Fig. 1D). Mass Spectrometry corroborated the enrichment of most α- and β-tubulin isotypes in TTC5 KO spheroids, albeit to a varying extent (Fig. 1E), strengthening the hypothesis that tubulin autoregulation is required to maintain tubulin quantity.

To investigate whether this increase in tubulin quantity affects microtubule intrinsic properties, we analyzed microtubule dynamics in 5-day-old spheroids transiently expressing comparable levels of GFP-tagged End-binding protein 3 (EB3-EGFP, Extended Data Fig. 1E). While microtubule growth lifetime remained unchanged (Extended Data Fig. 1F), EB3 comet analysis revealed 17% reduction in microtubule growth speed (16.1 μm/min versus 13.3 μm/min) (Fig. 1F-G) and 18% shorter growth length (Extended Data Fig. 1G) (1.6 μm versus 1.3 μm) in TTC5-deficient cells, indicative of altered microtubule stability.

Since modified dynamics are often linked to network rearrangement, we examined microtubule distribution in spheroids. Consistently, confocal imaging showed a redistribution of the microtubule density towards the cortex of TTC5 KO cells (Fig. 1H). Moreover, immunostaining for acetylated α-tubulin, a tubulin post-translational modification associated with increased microtubule stability^38^, confirmed an enrichment in tubulin acetylation in the absence of tubulin autoregulation (Fig. 1H-I, Extended Data Fig. 1H), suggesting an increase in microtubule stability upon loss of TTC5.

These findings establish that tubulin autoregulation is critical to maintain αβ-tubulin concentration and microtubule dynamics. The selective increase in tubulin mRNA and protein levels observed in both TTC5-deficient and -incompetent mutant spheroids confirms that these changes arise specifically from impaired tubulin autoregulation.

### Loss of tubulin autoregulation disrupts spheroid architecture and compromises cell viability

To investigate the broader consequences of impaired microtubule dynamics, we monitored spheroid growth from days 2 to 10 in parental cells and cells deficient in tubulin autoregulation (Fig. 2A). Using a trainable segmentation tool (see Methods for details on the quantification), we divided each spheroid into two regions, a dense central core and a less dense outer layer composed of proliferating cells (Fig. 2A ‘segmentation’). Over time, control spheroids retained a prominent core surrounded by a thin outer layer. In contrast, TTC5 KO spheroids exhibited progressive expansion, forming a large area of dispersed cells surrounding the spheroid, which we termed corona. This phenotype was rescued by TTC5^WT^ re-expression (Fig. 2A-B). By day 10, the corona area in the absence of TTC5 equaled or exceeded the size of the core, prompting us to perform most phenotypic analyses at this later time point. Notably, TTC5 KO spheroids re-expressing tubulin autoregulation-incompetent mutants phenocopied the knockout (Fig. 2A-B and Extended Data Fig. 2A). Similar architectural abnormalities were reproduced in an independent TTC5 KO clone (Extended Data Fig. 2B-C), strongly implicating loss of tubulin autoregulation as the cause of these defects.

**Fig. 2.**
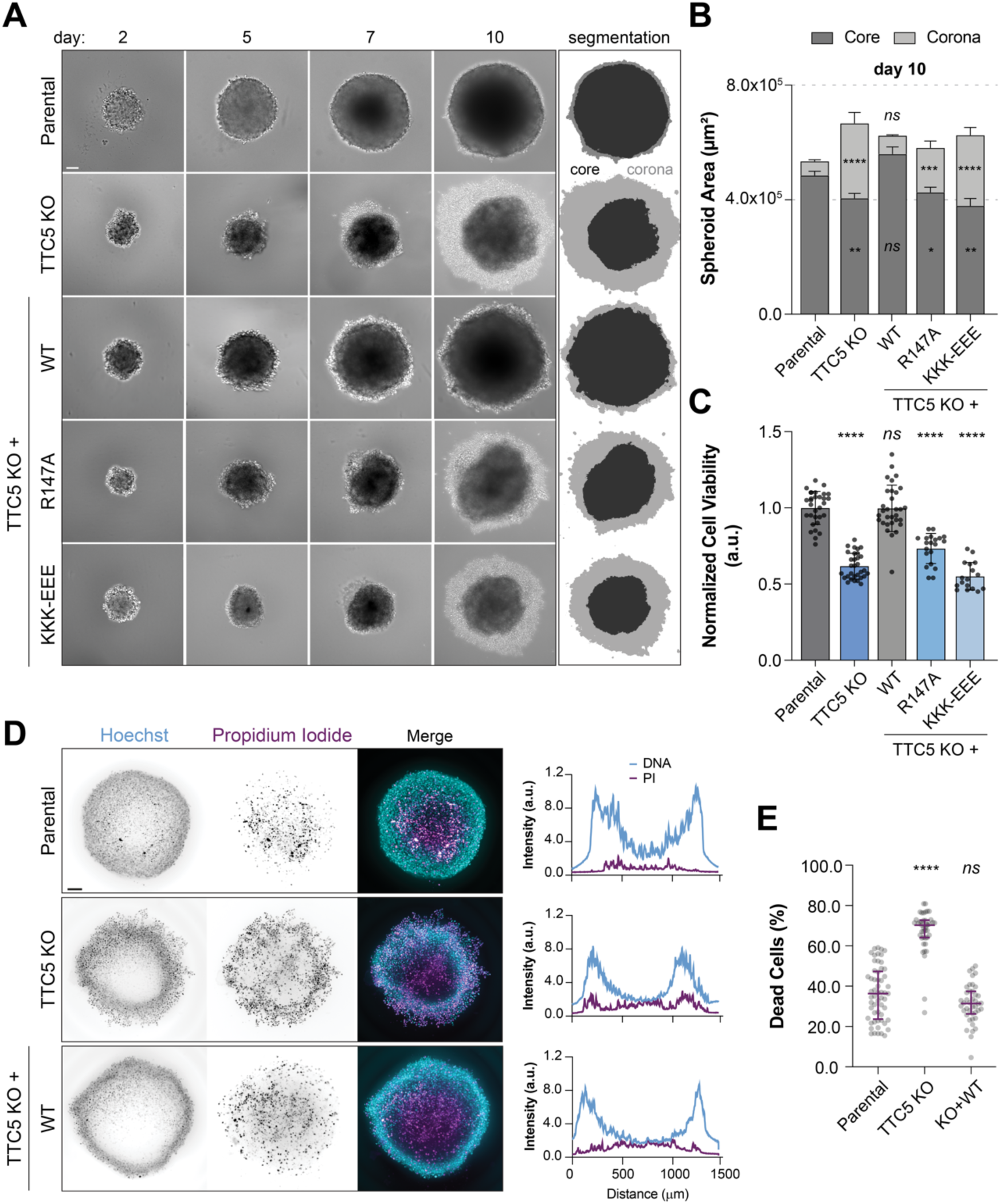
Tubulin autoregulation is required for normal spheroid growth and viability. (A) Representative brightfield images of spheroids from HeLa parental, TTC5 KO and the indicated Flag-TTC5 rescue cell lines (day 2 to day 10). Scale bar, 100 μm. Trainable Weka Segmentation masks are shown: core (dark grey), corona (light grey). (B) Spheroid core and corona areas measured at day 10. Data indicates mean ± SEM from minimum three biological replicates. *p* values indicate unpaired, two-tailed Student’s t-test relative to the parental cell line. (C) Cell viability determined by ATP content across the indicated cell lines, normalized to parental spheroids. Data show mean ± SD from three independent experiments. *p* values indicate unpaired, two-tailed Student’s t-test relative to the parental cell line. (D) Representative stills of HeLa parental, TTC5 KO and TTC5^WT^ rescue spheroid labelled with Hoechst (all nuclei, cyan) and Propidium Iodide (PI, dead cells, magenta). Scale bar, 100 μm. Graphs on the right show line scans measurements highlighting the distribution of all nuclei (blue) and dead cells (magenta) across the spheroid diameter. (E) Percentage of dead cells (PI-positive) from all segmented nuclei measured at day 10 in HeLa parental, TTC5 KO and TTC5^WT^ spheroids. Data are median ± interquartile range from a minimum of three independent experiments. *p* values from unpaired, Mann-Whitney test comparing ranks between TTC5 KO and TTC5^WT^ to parental spheroids. *\*\*\*\*p<*0.0001, *\*\*\*p<*0.001, *\*\*p<*0.01, *\*p*<0.05, *ns* - not significant.

To assess the impact on cell viability, we monitored ATP levels—as proxy of active cellular metabolism and survival—and observed a 40-50% reduction in TTC5 KO and autoregulation-incompetent mutant spheroids compared to controls (Fig. 2C). Next, we stained spheroids with Hoechst (labelling all nuclei) and Propidium Iodide (PI, to identify dead cells) to gain spatial insight into the distribution of viable and non-viable cells. In control and TTC5^WT^ spheroids, dead cells were mostly confined to the spheroid center (Fig. 2D), consistent with the expected hypoxic and nutrient-limited conditions. In contrast, TTC5-deficient spheroids displayed dispersed dead cells across the spheroid, particularly enriched at the corona (Fig. 2D). Quantification of PI-positive cells revealed a 2-fold increase in the percentage of dead cells in TTC5 KO spheroids compared to parental and TTC5^WT^ spheroids (Fig. 2E).

Halted progression through the cell-cycle may contribute to increased cell death^39^. To test this hypothesis, we analyzed DNA content by flow cytometry. This analysis revealed no significant changes across the different cell lines (Extended Data Fig. 3A), consistent with prior reports showing that TTC5-deficient cells complete mitosis within a similar timeframe as parental cells^18^. Supporting these observations, pharmacological inhibition of mitotic entry using the Cdk1 inhibitor Ro-3306 did not prevent corona expansion in TTC5 KO spheroids (Extended Data Fig. 3B-C).

One possibility was that peripheral cell death in autoregulation-deficient spheroids resulted from passive necrotic core disintegration. To test this hypothesis, we tracked individual living cells using Hoechst and PI. Time-lapse imaging showed that cell detachment and accumulation at the spheroid corona preceded cell death (Extended Data Fig. 3D, yellow arrowhead), implicating defective adhesion, rather than necrotic core dispersion, as a likely driver of the observed phenotype. In addition, a spheroid migration assay revealed no evidence of enhanced migration capacity in TTC5-deficient cells (Extended Data Fig. 3E-G).

Together, these data indicate that loss of tubulin autoregulation compromises multicellular architecture and cell viability not through altered proliferation or invasion, but possibly via impaired cell adhesion.

### Tubulin autoregulation ensures proper cell-cell and cell-ECM adhesion in 3D assemblies

Cell adhesion is essential for the organization of individual cells into 3D assemblies^40^. Microtubules play a key role in this process by delivering regulatory factors, scaffolding signaling molecules, and directly interacting with adhesion components, thereby coordinating cell-cell and cell-matrix adhesion^41^. We hypothesize that disruption in tubulin levels and microtubule dynamics may impair adhesion, potentially explaining the architectural defects observed in tubulin autoregulation-deficient spheroids. To test this hypothesis, we stained for bona fide adhesion proteins: N-cadherin, integrin α5, and fibronectin. Quantification of cortex-to-cytosol fluorescence ratios (Extended Data Fig. 4A; see Methods section for a full description) showed reduced cortical localization of adhesion proteins in TTC5 KO and mutant spheroids (Fig. 3A-D, Extended Data Fig. 4B-E). Notably, autoregulation-deficient cells (TTC5 KO, TTC5^R147A^ and TTC5^KKK-EEE^) exhibited more round shapes (aspect ratio, cell major axis/minor axis ratio, ∼1) than parental or TTC5^WT^ cells, consistent with loss of anchorage and weakened adhesion (Fig. 3E). Importantly, these adhesion proteins do not contain the TTC5-recognition motif and are therefore unlikely to be directly cotranslationally regulated by TTC5 (Extended Data Fig. 4F), implying that the observed effects are downstream consequences of disrupted tubulin homeostasis.

**Fig. 3.**
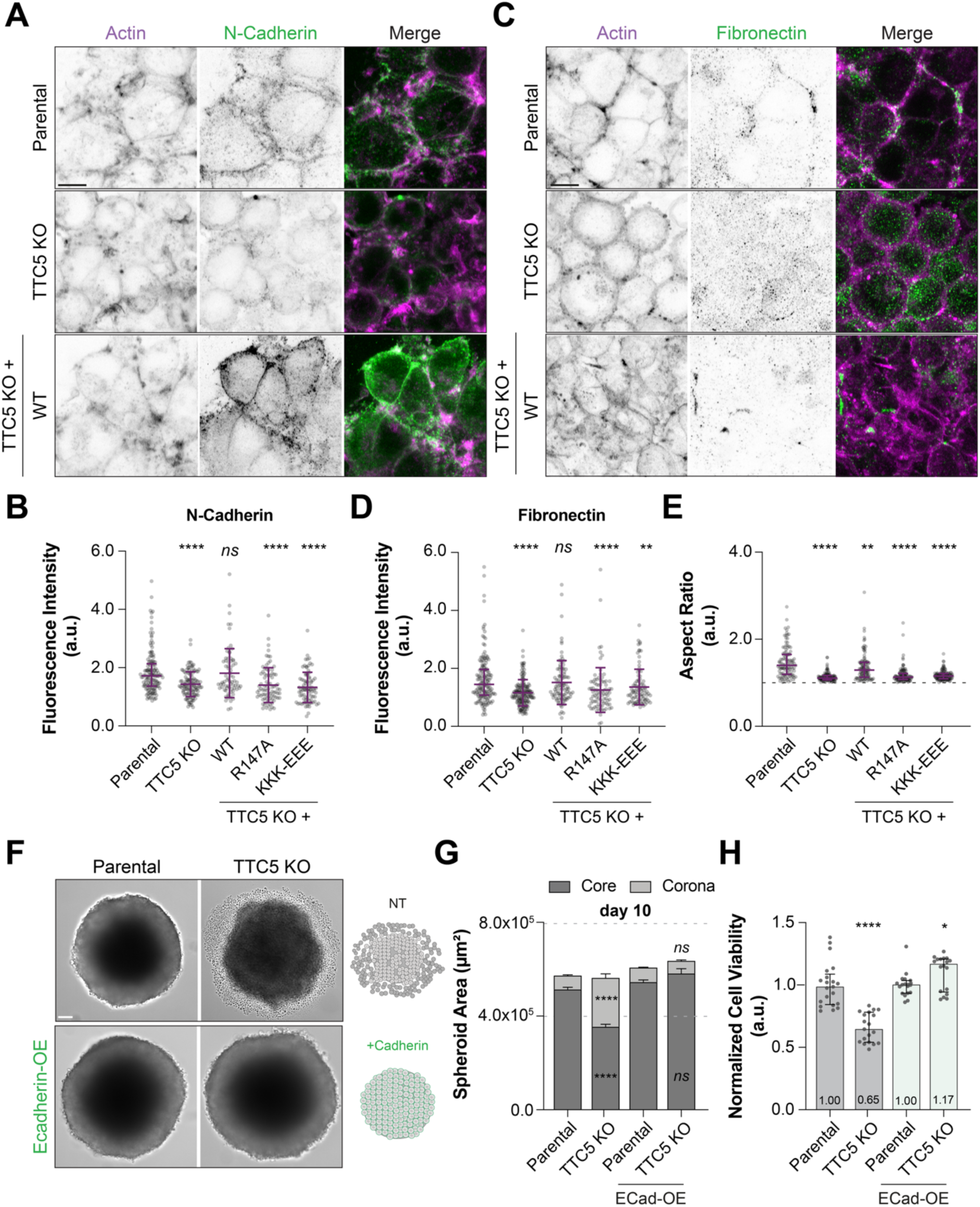
Tubulin autoregulation ensures proper cell–cell and cell–ECM adhesion in 3D assemblies. (A) Representative confocal images of 10-day HeLa parental, TTC5 KO and TTC5^WT^ spheroids cultured immunostained for N-Cadherin (green) and counterstained with Phalloidin-647 (magenta, Actin). Scale bar, 5 μm. (B) Quantification of N-Cadherin fluorescence intensity (cortex-to-cytoplasm ratio, see Extended Data Fig. 4A). Data are median ± interquartile range from two independent experiments. *p* values indicate unpaired Mann-Whitney test comparing ranks for each of the indicated cell lines with the parental cell line as reference. (C) Confocal images of spheroids stained for Fibronectin (green) and Phalloidin-647 (magenta, Actin). Scale bar, 5 μm. (D) Fluorescence intensity of Fibronectin (cortex-to-cytoplasm ratio). Data show median ± interquartile range from two independent experiments. *p* values indicate unpaired Mann-Whitney test comparing ranks for each of the indicated cell lines with the parental cell line as reference. Representative immunostainings of TTC5^R147A^ and TTC5^KKK-EEE^ mutants are shown in Extended Data Fig. 4B-C. (E) Aspect ratio measurements in HeLa parental, TTC5 KO and the indicated Flag-TTC5 rescue cell lines. Data are median ± interquartile range from three independent experiments. *p* values indicate unpaired Mann-Whitney test comparing ranks for each of the indicated cell lines with the parental cell line as reference. (F) Brightfield images of 10-day wildtype and E-Cadherin overexpressing HeLa parental and TTC5 KO spheroids. Scale bar, 100 μm. (B) Spheroid core and corona area quantification at day 10. Mean ± SEM from two independent experiments. *p* values indicate unpaired, two-tailed Student’s t-test comparing TTC5 KO spheroids to parental. (H) Cell viability determined by ATP content across the indicated cell lines, normalized to parental spheroids per condition. Data show median ± interquartile range from two independent experiments. *p* values indicate unpaired Mann-Whitney test comparing ranks for each group to the respective parental control. *\*\*\*\*p<*0.0001, *\*\*p<*0.01, *\*p<*0.05, *ns* - not significant.

In support of this hypothesis, overexpression of E-Cadherin in HeLa TTC5 KO spheroids, fully restored spheroid architecture and viability, and suppressed corona expansion (Fig. 3F-H). The same trend was observed when supplementing the culture medium with Matrigel, a basement membrane matrix rich in ECM components (Extended Data Fig. 5A-C). Consistently, co-culturing of TTC5 KO with parental cells restored spheroid morphology (Extended Data Fig. 5D) when parental cells constituted as little as 25% of the total cell population, suggesting that there is a collective input to overall spheroid adhesion.

Together, these findings establish that tubulin autoregulation is critical for maintaining proper cell-cell and cell cell-ECM adhesion, thus contributing to the architecture of multicellular 3D assemblies.

### Microtubule dynamics downstream of tubulin autoregulation control adhesion and tissue integrity

To further probe the functional link between tubulin autoregulation and spheroid architecture, we directly manipulated tubulin levels and microtubule dynamics and assessed their impact on spheroid morphology. First, we treated spheroids with low doses of T007-1, a small-molecule that covalently modifies Cys-239 on β-tubulin and triggers proteasomal degradation of both α- and β-tubulins^42^. While no effect was observed in parental spheroids, in TTC5 KO spheroids, T007-1 treatment led to a reduction in corona area rendering them indistinguishable from parental spheroids (Fig. 4A-B). This indicates that targeting the excess of tubulin accumulated in the absence of tubulin autoregulation can restore normal spheroid architecture.

**Fig. 4.**
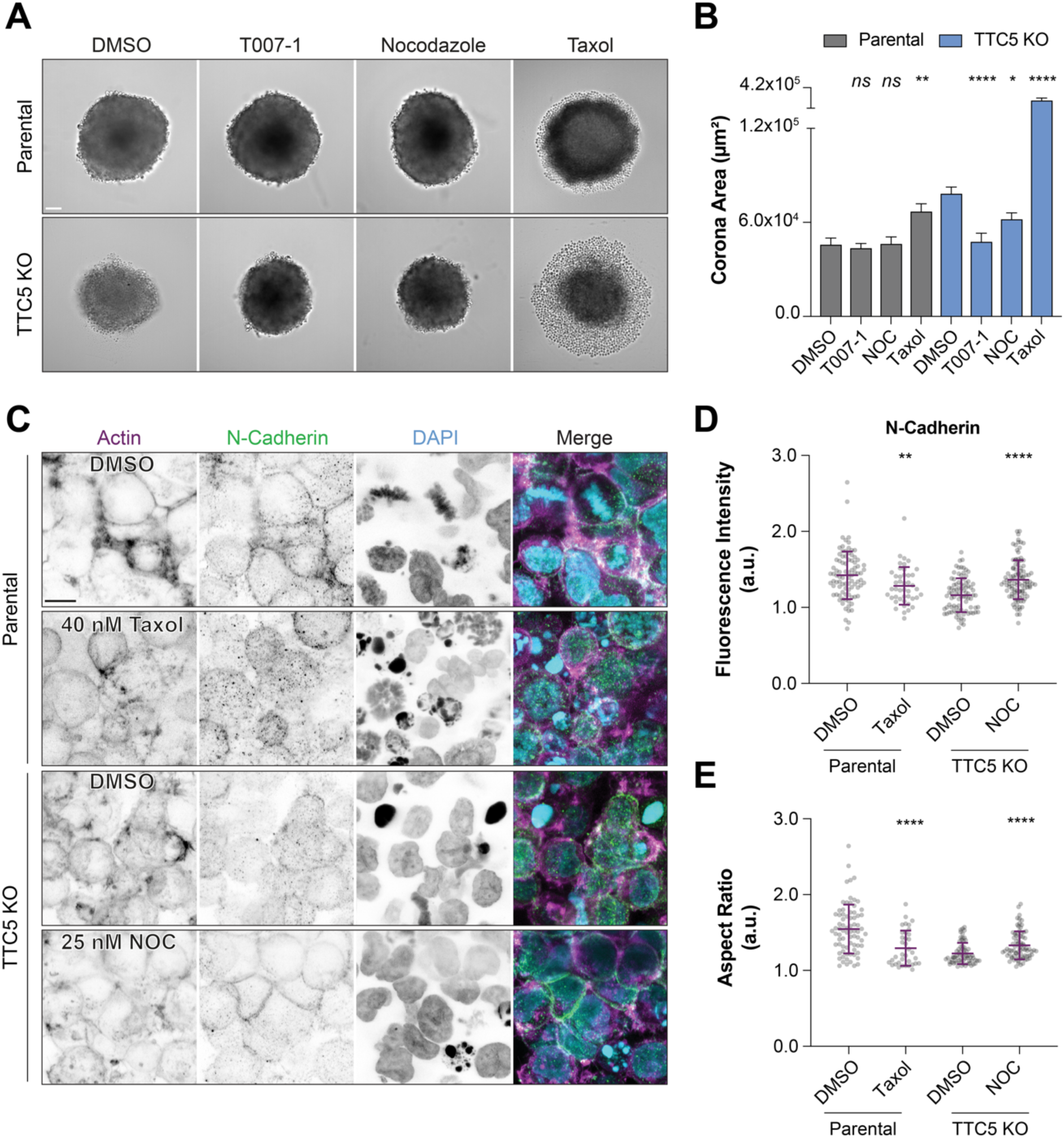
Microtubule dynamics downstream of tubulin autoregulation control adhesion and tissue integrity. (A) Brightfield images of 7-day HeLa parental and TTC5 KO spheroids treated for 48 hours with 1% DMSO, tubulin degrader T007-1 (400 nM), Nocodazole (NOC, 25 nM) or taxol (40/20 nM, respectively). Scale bar, 100 μm. (B) Spheroid corona areas at day 7. Data show mean ± SEM from three independent experiments. *p* values reflect unpaired, two-tailed Student’s t-test with DMSO condition as reference. (C) Confocal images of 7-day HeLa parental and TTC5 KO spheroids post-treatment, stained for N-Cadherin (green), Phalloidin-647 (magenta, Actin) and DAPI (cyan, DNA). Scale bar, 5 μm. (D) Cortex-to-cytoplasm N-Cadherin fluorescence intensity, while indicating median ± interquartile range from three independent experiments*. p* values indicate unpaired Mann-Whitney test comparing ranks for each of the indicated cell lines with the DMSO treatment as reference. (E) Aspect ratio of cells from parental and TTC5 KO spheroids upon the indicated treatments. Data show median ± interquartile range from three independent experiments. *p* values indicate unpaired Mann-Whitney test comparing ranks for each of the indicated cell lines with the DMSO treatment as reference. *\*\*\*\*p*<0.0001, *\*\*p<*0.01, *\*p<*0.05, *ns* - not significant.

Inducing mild microtubule destabilization with low-dose nocodazole (25 nM) reduced the corona area by ∼20% in TTC5-deficient spheroids. On the contrary, treatment with low-dose taxol, a microtubule stabilizing agent, led to the formation of the corona in parental spheroids, and exacerbated the phenotype in TTC5 KO spheroids, significantly increasing corona area (Fig. 4A-B, Extended Data Fig. 6A-B). Notably, TTC5 KO cells were particularly sensitive to taxol, suggesting that excessive microtubule stabilization has detrimental effects on spheroid architecture.

Restoring microtubule dynamics in TTC5 KO spheroids (low-dose nocodazole) reinstated N-Cadherin cortical localization and partly rescued cell shape (Fig. 4C-E). In contrast, 40 nM Taxol treatment impaired N-Cadherin localization and promoted a more rounded cell morphology in parental spheroids (Fig. 4C-E), thus mimicking the phenotype found in tubulin autoregulation-deficient cells.

These results highlight that microtubule dynamics downstream of tubulin autoregulation are key for the localization of adhesion molecules. Imbalances in microtubule stability profoundly disrupt cell adhesion and 3D tissue organization, underscoring the importance of finely tuning tubulin homeostasis for maintaining tissue integrity and viability.

## Discussion

Tubulin autoregulation is a conserved feedback mechanism that maintains balanced tubulin biosynthesis by selectively degrading tubulin-encoding mRNAs when αβ-tubulin levels are excessive^43^. Recent mechanistic studies established that, when liberated from soluble αβ-tubulins, TTC5 recognizes nascent α- and β-tubulin chains during translation and—via recruitment of SCAPER and the CCR4-NOT deadenylase complex—initiates mRNA deadenylation and decay^18–20^. These discoveries have enabled the development of the first molecular tools to manipulate tubulin autoregulation in cells, opening the door to functional analyses of this pathway.

Given its function in limiting runaway tubulin biosynthesis, tubulin autoregulation has been assumed to act as a quantity control pathway. Yet, pioneering studies relied exclusively on microtubule-targeting agents to trigger tubulin autoregulation, leaving its physiological relevance unexplored. In this study, we uncover a critical role for TTC5-mediated tubulin autoregulation in maintaining tubulin abundance, regulating microtubule dynamics, and preserving spheroid architecture. Combining genetic, biochemical, and imaging approaches, we demonstrate that loss of tubulin autoregulation leads to increased tubulin levels, disrupted microtubule dynamics and weakened cell adhesion, therefore compromising spheroid morphology and viability. This work establishes tubulin quantity control as a core regulator of multicellular homeostasis.

We propose a model in which TTC5-mediated response to excess unpolymerized tubulin couples translation to mRNA degradation, limiting mRNA accumulation and preventing tubulin buildup. At first glance, our findings seem at odds with previous work reporting moderately increased tubulin mRNA, yet unchanged protein levels upon loss of tubulin autoregulation in monolayer cultures^18^. However, monolayer systems may lack physiological signals that activate autoregulation, displaying minimal pathway activity at steady state. In contrast, 3D cultures such as spheroids exhibit higher autoregulation activity, allowing functional consequences to emerge. Consistent with this model, a switch from 2D to 3D culture model was previously associated with differential tubulin gene expression^27^. Together, these observations suggest that tubulin autoregulation is highly context dependent. Future work exploring how extracellular cues, such as ECM composition, matrix stiffness and tissue geometry, curb autoregulation activity will help to elucidate the mechanism that tunes tubulin output in response to environmental demands.

The relationship between tubulin concentration and microtubule dynamics is well established *in vitro*, where polymerization rate and dynamic instability scale linearly with tubulin availability^8–11^. In cells, however, this link has remained elusive due to lack of tools to selectively manipulate tubulin levels. Computational models have attempted to address this gap, but generally fall short due to the large complexity of the tubulin network and vast microtubule regulators^44–46^. Manipulating tubulin autoregulation represents a unique molecular handle to upregulate tubulin levels in cells. Leveraging this strategy, we reveal that even subtle changes in tubulin abundance considerably reduce microtubule growth rate and growth length in cells. These findings may explain previously reported loss of mitotic fidelity in tubulin autoregulation-deficient cells^18–20^.

Curiously, similar changes in dynamics have been observed upon loss of core microtubule-associated proteins, such as end-binding protein 3 (EB3)^47,48^, CAP-Gly Domain Containing Linker Protein 1 (CLIP1)^49^, or Cytoplasmic Linker Associated Protein 1 (CLASP1)^50^, placing tubulin levels on par with classical binding proteins in shaping microtubule behavior. Future studies using tubulin small molecule degraders^42,51^ to profile microtubule dynamics in regimes when tubulin building blocks are depleted could complement our approach, and offer a comprehensive view on how microtubule dynamics scale with tubulin quantity in cells.

Tightly regulated dynamics are critical for proper microtubule network organization and function. Consistent with this, tubulin autoregulation-deficient cells exhibit reorganization of the microtubule network, with acetylated microtubules enriched at the cell cortex. Such architectural change is likely to disrupt directional transport and the delivery of key adhesion molecules to the plasma membrane, contributing to reduced cortical localization of cadherins and integrins^52–55^. Defective trafficking—perhaps due to altered kinesin or dynein motility along stabilized tracks—may underlie the failure to establish and maintain robust intercellular and ECM adhesions, and ultimately compromise the integrity of multicellular assemblies such as spheroids^41^. Endogenous labeling of the relevant ECM proteins and development of sophisticated imaging approaches adapted to 3D culture models will be required to test this hypothesis.

Our study provides a proof-of-principle that tubulin autoregulation is required for the integrity of multicellular assemblies. While we focused on tumor-like spheres, this pathway may contribute to the organization of diverse tissues, both normal and pathological. Although tubulin autoregulation appears functional across diverse human cell types^56–58^, it is likely that different tissues require distinct tubulin levels to support specialized functions. Such specialization could be achieved through additional upstream regulators of TTC5 and/or SCAPER and fine-tuning of the tubulin autoregulation pathway. Similarly, tubulin autoregulation may act in pathological conditions, as seen in heart hypertrophy where it seems to rewrite the tubulin isotype code^59^. The development of *in vivo* models and reporters of tubulin autoregulation will be key to charting this pathway’s activity across biological contexts. Mutations in TTC5 and SCAPER have been linked to a range of human diseases, including hereditary dystonia, neurodevelopmental syndromes, and various cancers^23,60,61^, though the molecular underpinnings remain unknown. Our work raises the possibility that these phenotypes stem from disrupted microtubule behavior and function due to impaired tubulin quantity control—an idea that warrants further mechanistic study.

Given the importance of microtubule dynamics in cancer, and the observation that tubulins are often upregulated in tumors^62^, targeting TTC5 or the broader autoregulation machinery may represent a novel therapeutic strategy. Supporting this hypothesis, our data reveal increased sensitivity of tubulin autoregulation-deficient spheroids to microtubule-stabilizing poisons such as paclitaxel. Future work is required to profile the druggability of the tubulin autoregulation pathway and evaluate its potential for tumor-selective intervention.

Together, our findings place tubulin autoregulation at the core of microtubule regulation and multicellular integrity, offering new insight into how cells orchestrate cytoskeletal dynamics via regulated protein biosynthesis. By uncovering a physiological role for this long-elusive feedback loop, we lay the groundwork for deeper exploration of protein quantity control and its implications in health and disease.

## Methods

### Cell Culture and Cell Lines Generation

Flp-In T-REx HeLa cells (*Thermo Fisher Scientific, #R71407*) were maintained in DMEM with GlutaMAX (*Thermo Fisher Scientific, #10566016*) supplemented with 10% Fetal Bovine Serum (*FBS, Pan Biotech, #P30-3306*) and 1x Penicillin-Streptomycin (*Thermo Fisher Scientific, #15140122*) at 37°C in humidified conditions with 5% CO_2_.

Stable cell lines expressing TTC5 wild-type (WT), and autoregulation-incompetent mutants were generated previously^18^. E-Cadherin (*Addgene, #28009*) was subcloned into a pcDNA5/FRT/TO-EGFP vector. Flp-In T-REx HeLa parental and TTC5 KO cells were co-transfected with pOG44 (Flp recombinase) and the plasmid of interest at a 1:1 ratio using Lipofectamine 3000 (*Thermo Fisher Scientific, #L3000015*), following the manufacturer’s protocol (*#K650001*). The following day, cells were selected with 0.2 mg/mL Hygromycin B (*Corning, #30-240-CR*) and 10 μg/mL Blasticidine S (*Thermo Fisher Scientific, #R21001*) for 10-15 days. Transgene expression was induced with 200 ng/mL Doxycycline (*Sigma-Aldrich, #D9891-1G)* for a minimum of 24 hours. All cell lines were routinely checked for mycoplasma contamination.

For transient expression of EB3-EGFP, ∼400.000 cells (parental and TTC5 KO) were seeded in 6-well plates and transfected with 0.5 μg of EB3-EGFP plasmid (kindly provided by Prof. Charlotte Aumeier, University of Geneva) using 1:5 ratio of DNA:Lipofectamine 3000. After 24 hours, cells were trypsinized and grown in spheroids for 5 days.

### 3D Cell Culture (Human Spheroids)

Spheroids were generated by seeding 1,000 cells Flp-In T-REx HeLa cells per well into Nunclon Sphera 96-well U-shaped-bottom plates (*Thermo Fisher Scientific #174925)* in complete growth medium containing 200 ng/mL Doxycycline. Plates were centrifuged at 290 × g for 5 minutes at room-temperature and incubated at 37 °C in a humidified 5% CO_2_ atmosphere. Spheroids were maintained for up to 10 days in culture, with two media changes performed between days 5 and 10.

For rescue experiments, Matrigel (*Corning, #356231, Lot n° 2262003*) was added to the medium at 1 mg/mL on either day 5 or day 7, with analysis of spheroid growth and viability on day 10.

For drug treatment assays, microtubule-targeting agents and Cdk1 inhibitor were diluted in complete medium and added at day 5 or day 7. Final concentrations used were: Nocodazole 25nM, Paclitaxel 20, 40 or 80 nM, T007-1 400 nM, Colchicine 5 μM, and Ro-3306 5 μM at the indicated treatment duration.

### mRNA quantification by RT-qPCR

Parental, TTC5 KO, and indicated rescue cell lines were cultured as spheroids with 200 ng/mL Doxycycline for 10 days. For tubulin autoregulation assay, spheroids were treated with 5 μM Colchicine (*Sigma-Aldrich, #PRH1764*) or 0.01% DMSO (*Sigma-Aldrich, #D2438)* for 6 hours. Spheroids were dissociated with Accutase (*Thermo Fisher Scientific, #00-4555-56*), and RNA extracted using the NucleoSpin RNA Mini Kit (*Macherey-Nagel, #740955*). DNA synthesis was performed from 1 μg RNA using the SensiFAST cDNA Synthesis Kit (*Bioline, #BIO-65054*), as per manufacturer’s instructions. qPCR was carried out adding 10 ng of cDNA, 2x PowerUp SYBR Green Master Mix (*Thermo Fisher Scientific, #A25777*), and indicated primers on a Bio-Rad thermocycler.

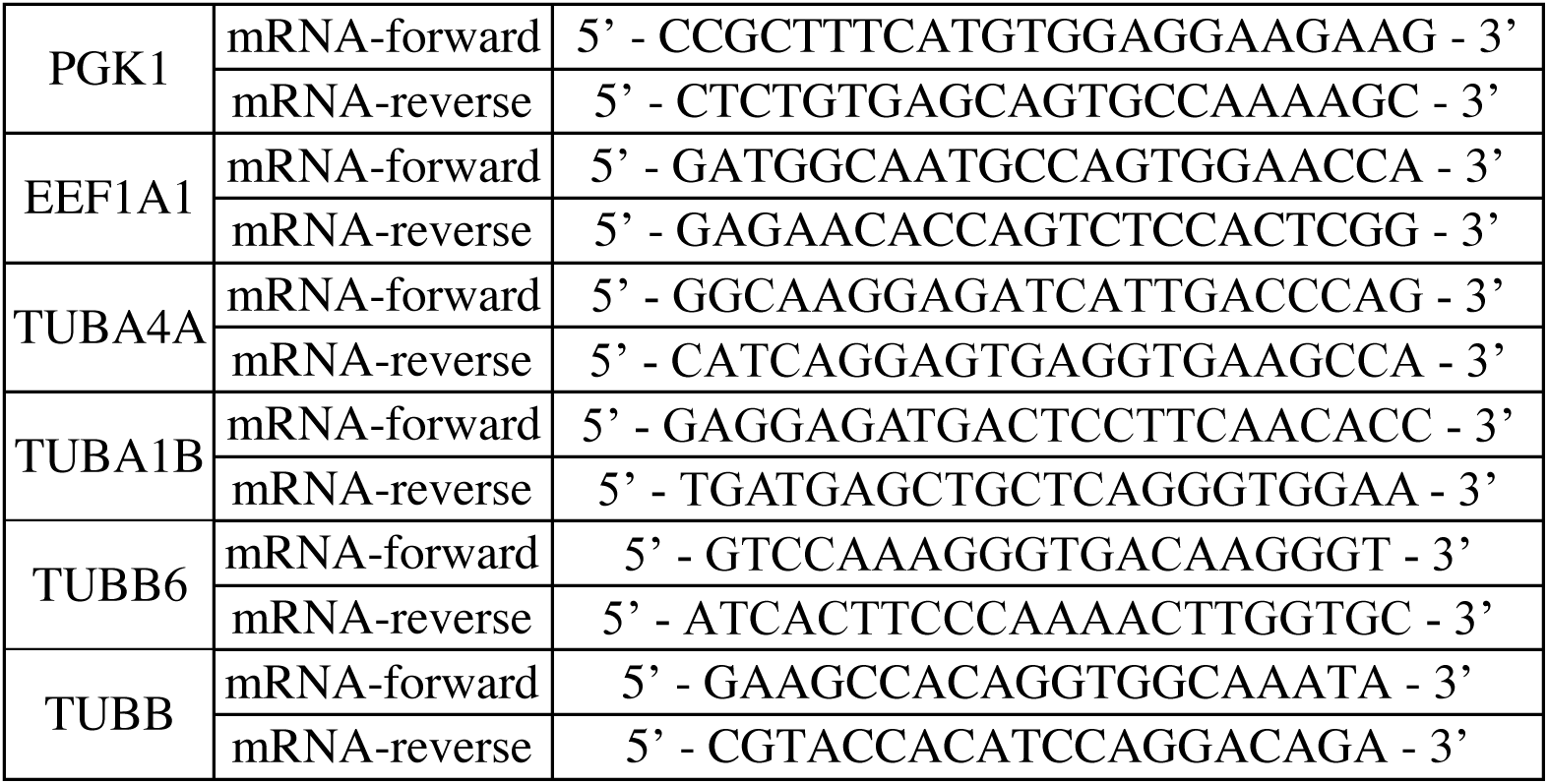

Relative expression was analyzed via the ddCt method^63^, normalized to reference housekeeping genes (PGK1 and EEF1A1), and expressed relative to DMSO or parental controls. Experiments include a minimum of three biological replicates. Data analysis and plotting were conducted using R/RStudio (v2022.12.0+353).

### Western blotting and Quantification

Spheroids were dissociated into single cells with Accutase and lysed in RIPA buffer (50 mM Tris-HCl, pH 8.0, 150 mM NaCl, 1% Triton X-100, 0.5% sodium deoxycholate, 0.1% SDS) supplemented with protease (1:100, Halt™ *Thermo Fisher Scientific, #78429*) and phosphatase inhibitors (1:25, PhosSTOP, *Roche, #4906845001*).

Protein lysates were separated via SDS-PAGE on 4-12% Tris-Glycine pre-cast gels (*Thermo Fisher Scientific, #XP04125BOX*) or 10% homemade Tris-Glycine gels, and transferred to 0.2 μm nitrocellulose membranes (*Amersham, Cytiva*). After Ponceau S staining (*Roth, #5938.1)*, membranes were blocked in 5% non-fat dry milk prepared in PBS containing 0.2% Tween-20 (*Santa Cruz, #sc-29113*). Primary antibodies: anti-α-tubulin (1:5000, *Proteintech, #11224-1-AP, Lot n° 0013194*) and anti-GAPDH antibody (1:10000, *Cell Signaling Technology, #2118S, Lot n° 10*) were incubated overnight at 4°C. Detection was carried out using IRDye-conjugated secondary antibodies LI-COR 550 (1:10000, *Azure Biosystems, #AC2159, Lot n° 240322-56*) and LI-COR 800 (1:5000, *Thermo Fisher Scientific, #A32735, Lor n° WA318266*) and imaging was performed with the Sapphire system (*Azure Biosystems*).

Band intensities were quantified in Fiji (*ImageJ, 1.54f*). Briefly, rectangular regions of interest were manually drawn around each band, keeping the area constant throughout all lanes. Upon background subtraction, mean intensity values of α-tubulin were normalized to the corresponding GAPDH signal. Data were expressed as α-tubulin/GAPDH ratios.

### Sample preparation for proteomic analysis

Spheroids from parental and TTC5 KO HeLa cells (10 days in culture) were harvested in PBS. Samples were prepared for liquid chromatography/mass spectrometry (LC/MS) using the phase-transfer surfactant method, with minor modifications. First, proteins were extracted and solubilized using buffer containing 12 mM sodium deoxycholate, 12 mM sodium N-dodecanoylsarcosinate, and 100 mM Tris pH 9.0, with EDTA-free Protease Inhibitor Cocktail (*Roche*). Samples were sonicated for 4 minutes using a Bandelin Sonorex ultrasonic bath (*FAUST*) with 20-seconds on/20-seconds off cycles. Cell debris was removed after centrifugation at 18,000 × g for 20 minutes at 4 °C. Samples were reduced with 10 mM TCEP at 37 °C for 30 minutes and alkylated with 20 mM iodoacetamide in the dark at room temperature for 30 minutes. Alkylation reactions were quenched with 75 mM cysteine at room temperature for 10 minutes. Samples were diluted with 3.1 volumes of 50 mM ammonium bicarbonate. Lysyl endopeptidase and trypsin (*Promega*) were added at a 50:1 ratio of sample protein:enzyme (w/w) and samples were digested for 16 hours at 37 °C. Afterward, 1.77 volumes of ethyl acetate were added, and samples were acidified with trifluoroacetic acid (TFA), which was added to 0.46% (v/v). Following centrifugation at 12,000 × g for 5 minutes at room temperature, samples separated into two phases. The upper organic phase containing sodium deoxycholate was removed, and the lower aqueous phase containing digested tryptic peptides was dried using a centrifugal vacuum concentrator. Digested peptides were dissolved in 300 μL of 0.1% (v/v) TFA in 3% acetonitrile (v/v). Samples were sonicated for 1 minute, centrifuged at 15,000 × g for 15 minutes, and desalted using MonoSpin C18 columns (*GL Sciences Inc.*). Peptides were eluted from C18 columns using 0.1% TFA in 50% acetonitrile and dried in a vacuum concentrator. Tryptic peptides were dissolved in 0.1% (v/v) formic acid in 2% (v/v) acetonitrile for mass spectrometry analysis.

### Mass Spectrometry

Samples were measured on an Easy Nano LC - Orbitrap Fusion System equipped with a nanospray flex™ ion source (*Thermo Fisher Scientific*). Peptides were separated on a 1.9-μm particle, 75-μm inner diameter, 15 to 20-cm filling length homemade C18 column. A flow rate of 300 nL/min was used with a 114-minute gradient (2–25% solvent B in 100 minutes, 25– 45% solvent B in 7 minutes, 45–75% solvent B in 7 minutes). The gradient was followed with two rounds of washing steps, in each step, the gradient switched to 98% solvent B in 1 minute and remained for 2 ×, then switched to 2% solvent B in 1 minute and remained for 2 minutes. After the second round of washing, 2% solvent B was reached in 1 minute and remained for 14 minutes for system equilibration. Solvent A was 0.1% (v/v) formic acid in LC/MS grade water and solvent B was 0.1% (v/v) formic acid in 100% (v/v) acetonitrile. The ion source settings from Tune were used for the mass spectrometer ion source properties.

For data-independent acquisition (DIA), data were acquired with 1 full MS and 38 overlapping isolation windows constructed covering the precursor mass range of 350–1200 m/z. For full mass spectrometry, Orbitrap resolution was set to 120,000. AGC target was set to custom with normalized AGC target set to 250%, and maximum injection time (IT) was set to 60 milliseconds. DIA segments were acquired at resolution of 30,000. AGC target was set to custom with normalized AGC target set to 2000%, and a dynamic maximum IT. HCD fragmentation was set to normalized collision energy of 27%.

For protein identification and quantification raw files were analyzed using the directDIA workflow in Spectronaut software (*Biognosys*) with default settings. Identification was performed using the Uniprot reference proteome for human. Digestion enzyme specificity was set to Trypsin/P. Modification included carbamidomethylation of cysteine as a fixed modification, and oxidation of methionine and acetyl (protein N-terminus) as variable modifications. Up to two missed cleavages were allowed. FDR was estimated with the mProphet approach and set to 0.01 at both peptide precursor level and protein level in at least 80% of samples. For two-group comparison, differential abundance testing was performed with unpaired t-test. Q-values were the multiple testing corrected p-values. Data processing and visualization were done in R/RStudio (v2022.12.0+353).

### Microscopy-based assays

#### Scanning Confocal Microscopy (EB3 Comet Tracking)

Five-day spheroids transiently expressing EB3-EGFP were transferred to µ-Slide 15 Well 3D dishes (*iBidi, #81506*) and imaged in phenol-red-free Leibovitz’s L-15 medium (*Thermo Fisher Scientific, #21083027*) supplemented with 10% FBS. Imaging was performed on a temperature-controlled Zeiss LSM980 confocal microscope fitted on an inverted Axio Observer 7 Microscope with an Airyscan detector optimized for a Plan-ApoChromat 63x/1.40 NA oil objective (0.085 μm/pixel). Acquisition optics were composed of a laser line 488 nm with 5.5x digital zooming. Time-lapse imaging captured one plane every 1.5 seconds for 1 minute. ZEN (blue edition, 3.3.89.0008) software was used for image acquisition and Airyscan processing.

EB3 comets dynamics were analyzed using the multiple-particle tracking MATLAB software u-track^64^. Custom parameters were set for: (1) comet detection: difference of Gaussians filter parameters 1-4 pixels; watershed segmentation 5 standard deviations; (2) tracking: maximum gap to close 3 frames; minimum length of track segments 4 frames; search radius 2-8 pixels; maximum forward/backward angles 40°/10°; break non-linear tracks; and (3) microtubule dynamics classification. Due to the low frequency of shrinkage events, we focused our analysis on microtubule growth lifetime, length, and speed. Cytoplasmic EB3-EGFP intensities, measured as the mean intensity on a region of interest in the cytoplasm of each analyzed cell, were used to exclude outliers (average intensity ± standard deviation, grouped per experiment).

#### Scanning Confocal Microscopy (Fixed Spheroids)

After processing spheroids for immunofluorescence (see section below), imaging was performed on the same confocal setup. Excitation wavelengths were composed of 405 nm, 488 nm, 561 nm, and 647 nm laser lines, with 1.5-2x digital zooming, and z-step of 0.2 μm. Due to the large spheroid volume and constraints in imaging depth, acquisition was limited to the peripheral regions of the spheroids. All images show max-intensity projections (representation only).

For analysis, sum-projection images were obtained in Fiji. Cell cortex was manually defined (using the segmented line tool, 20 pixel) with Phalloidin/actin staining as reference. Acetylated-tubulin and α-tubulin fluorescence mean intensities were measured along the cortical region, and results presented as a ratio. Mitotic cells were excluded from the quantification. For aspect ratio measurements (defined by the ratio of the cell width to its heigh), polygon tool was used to mark the cell cortex, using Phalloidin/actin staining as reference.

Cortical and cytoplasmic intensities were extracted via line scans (20-30 pixel) placed from the center of individual cells to the cortex, using actin as a reference. Cortex-to-cytoplasm ratio of adhesion proteins was defined as *p/m*, where *p* is the mean intensity within ± 1 μm from peak intensity (cell cortex) and *m* is the mean intensity 1–2 μm from *p* (into the cytoplasm).

#### Widefield microscopy (Spheroid Growth and Drug Response)

Spheroid growth was monitored from day 0 to day 10 on a Nikon Eclipse Ti2-E inverted microscope (Nikon), equipped with a Kinetix sCMOS camera (Photometrics), Spectrax Chroma light engine for fluorescence illumination (Lumencor), and an incubation chamber with 37 °C with controlled humidity and 5% CO_2_ (OkoLab). Brightfield multiple stage positions were acquired using NIS Elements (Nikon) equipped with a Plan Apochromat Lambda 10x objective (NA 0.45, Nikon) sampling 0.64 μm/pixel. Single plain images or maximum intensity projections were prepared in Fiji and spheroid area analyzed using a manual or automatic pipeline that runs trainable Weka Segmentation plugin (version 3.3.2) to identify core, corona, and background classes. Binary images were generated from the probability maps using Otsu threshold (auto adjustment was used as default, but manual adjustments were required in some instances), and areas (core, corona) measured in microns. <u>Widefield microscopy</u> (Cell Viability assay)

Nine-day spheroids stained with Hoechst and PI to visualize all nuclei and dead cells, respectively, were imaged in steps of 5 μm, every 20 minutes for 10 hours. Images were denoised (NIS Elements), and maximum intensity projections of representative examples were prepared in Fiji and exported as still images.

#### High-throughput live-cell imaging (Cell Viability assay)

Ten-day spheroids were stained with Hoechst and PI and imaged using an ImageXpress Micro Confocal automated microscope (*Molecular Devices*™) equipped with a Plan Apochromat Lambda 10x objective (0.45 NA, Nikon). Image segmentation was performed using a custom module editor MetaXpress from Molecular Devices. Briefly, single nuclei were masked and segmented using Hoechst signal. Percentage of dead cells was calculated by dividing the number of PI positive cells by the total number of segmented nuclei.

#### High-throughput live-cell imaging (Spheroid migration assay)

Seven-day spheroids were imaged in a 384-well plate coated with Matrigel for 48 hours (see ‘Spheroid migration assay’ section) using a IXM confocal automatic microscope with the same specs as indicated above. Spheroid and migration areas were segmented and measured with MetaXpress Custom Module editor software using brightfield channel to segment the two objects.

### Immunofluorescence

Spheroids (7 or 10 days) were fixed in 4% paraformaldehyde (PFA, *Electron Microscopy Sciences #15713*) in PBS for 30 minutes at 37°C. Permeabilization was performed using PBS-0.2% Triton (*Sigma-Aldrich X-100, #9036-19-5*) for 30 minutes at room-temperature. Spheroids were washed 3x with PBS and incubated for 2 hours at room-temperature with blocking solution: 2% BSA (*PAN Biotech, #P06-1391500, Lot n° H210207*) diluted in PBS with 0.02% Tween 20 (PBS-T). Primary antibodies: anti-acetylated tubulin (1:200, *Sigma-Aldrich, #T7451, Lot n° 0000312701),* anti-α-tubulin (*1:200, Proteintech, #11224-1-AP, Lot n° 0013194),* anti-N-Cadherin (1:200, *Proteintech, #22018-1-AP, Lot n° 00111704*), anti-Fibronectin (1:1000, clone 1801, kind gift from Prof. Bernhard Wehrle-Haller, University of Geneva) and anti-Integrin α5 (1:10, AA430-M2a, kind gift from Prof. Bernhard Wehrle-Haller, University of Geneva) were diluted in blocking solution and incubated overnight at 4°C. Subsequently, spheroids were washed 3x with PBS-T and incubated for 2 hours at room temperature with the corresponding secondary antibody: Alexa 488 (1:500, *Thermo Fisher Scientific, #A32731, Lot n° WC318798*) and 555 (1:500, *Thermo Fisher Scientific, #A32727, Lot n° WA316324*), together with DAPI (1:5000, 5mg/mL*, Thermo Fisher Scientific, #D1306*) and Phalloiding-647 (1:250, *Thermo Fisher Scientific, #A22287, Lot n° 1583100*). After three washes with PBS-T, spheroids were kept in PBS at 4°C until imaging.

### Cell Viability Measurements

Ten-day spheroids were stained with PI (1:2000, 1 mg/mL, *Sigma-Aldrich, #81845*) and Hoechst 33342 (1:2000, 10 mg/mL, *Molecular Probes, #H-3570*) for 2 hours prior to imaging. Spheroids were processed for widefield, or high-throughput microscopy as indicated in the ‘Microscopy-based assays’ section.

For ATP-based viability assay, spheroids at day 7 and 10 were incubated with CellTiter-Glo 3D reagent (Promega, #G9681) for 30 minutes, following manufacturer’s instructions. Luminescence readout was measured on a Cytation 3 imaging reader (Gen5 3.14).

### Flow cytometry

Spheroids (10-day) were dissociated with Accutase for 20 minutes at 37°C. After centrifugation (300 × g for 5 minutes at room-temperature), cells were fixed in 90% cold methanol (-20 °C, overnight. Following washes with PBS, DNA was stained with PI (1:60, 1 mg/mL) together with RNase treatment (1:150, *Roche, #11119915001*). Cell cycle profiles, based on PI staining, were acquired on a Gallios cytometer (*Beckman Coulter*), and analyzed using Kaluza flow cytometry software (*Beckman Coulter)*.

### Spheroid migration assay

Seven-day spheroids were manually transferred to a 384-well plate (*Corning, #353962*) coated with 125 μg/mL Matrigel (*Corning, #356231*, 2 hours at room-temperature) following an established protocol^65^. Cell migration was recorded for two days (timepoints 0, 24 and 48 hours) using a IXM confocal automatic microscope (check ‘Microscopy-based assays’ section for imaging details).

### Statistical analysis

For all data, the number of independent biological replicas and statistical tests with *p* values are detailed in the figures and figure legends. No statistical method was used to predetermine the sample size. The experiments were not randomized. The investigators were not blinded to allocation during experiments and outcome assessment. Statistical analyses were performed in R and GraphPad Prism 8, which reports exact *p* values up to four decimal places, with smaller values shown as *p*<0.0001. D’Agostinho-Pearson omnibus normality test was used to determine if the data followed a normal distribution. If α = 0.05, a statistical significance of differences between the population distributions was determined by Student’s t test. If α < 0.05, statistical analysis was performed using a Mann-Whitney Rank Sum test. ROUT method (Q = 1%) was used to identify outliers in the WB quantification dataset. Plot in Extended Data Fig. 1D shows the cleaned data. For each graph, ∗∗∗∗*p*<0.0001, ∗∗∗*p* <0.001, ∗∗*p* <0.01, ∗*p*<0.05 and *ns* - not significant.

## Acknowledgments

We are grateful to P. Meraldi and H. Maiato for critical reading of the manuscript, and all members of the Gasic team for their valuable feedback and support. Reagents and helpful input were generously provided by C. Aumeier and B. Wehrle-Haller. We thank D. Moreau (ACCESS platform, University of Geneva) for assistance with high-throughput imaging and data analysis in spheroids; the Mass Spectrometry Core Facility at the University of Geneva for support with sample preparation and mass spectrometry analysis; and F. Lemaitre for helping to automate the Trainable Weka Segmentation in Fiji. This work was supported by the Swiss National Science Foundation (SNSF, grant PCEFP3_194312 to I.G.), the Republic and Canton of Geneva, Switzerland (DIP, to I.G.) and a postdoctoral fellowship from EMBO (ALTF 258-2023) awarded to A.C.A..

## Author contributions

A.C.A. performed all experiments and conducted the formal data analysis. C.L. contributed to western blotting, RNA quantification, RT-qPCR, and immunostaining experiments. A.C.A. conceptualized and validated the experiments/methodologies, prepared the figures, and wrote the initial draft of the manuscript. I.G. conceived the study, oversaw its design and implementation, provided supervision, and revised the manuscript. All authors contributed to manuscript editing.

---

**Extended Data Fig. 1.**
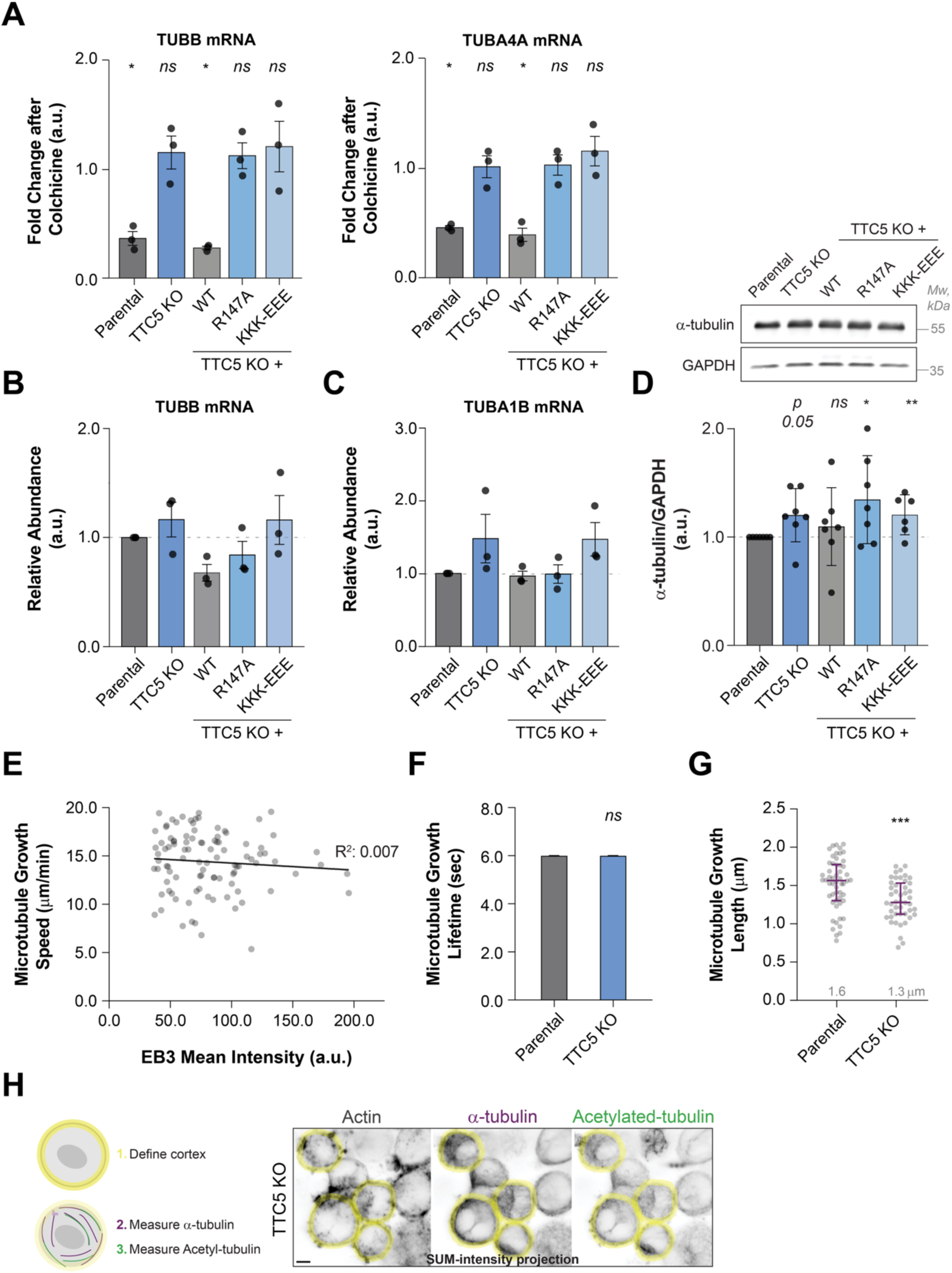
(A) Autoregulation assay with HeLa parental, TTC5 knockout, and the indicated Flag-TTC5 rescue cell lines, normalized to DMSO condition. Data represent mean ± SD TUBB and TUBA4A mRNA levels after 6 hours colchicine treatment from three independent experiments. *p* values indicate unpaired, two-tailed Student’s t-test for each of the cell lines with DMSO as reference. (B) Relative abundance of TUBB and (C) TUBA1B mRNA in HeLa parental, TTC5 knockout, and the indicated Flag-TTC5 cell lines in 10-days old spheroids, normalized to housekeeping transcripts and the parental levels. Data show mean ± SD from three independent experiments. No significant differences were detected using unpaired, two-tailed Student’s t-test for each of the indicated cell lines compared to parental cell line. (D) Western blot analysis of total α-tubulin protein levels in the indicated cell lines. Quantification shows α-tubulin intensity normalized to loading control (GAPDH) and parental cell line. Data show mean ± SD from five independent experiments (including technical replicates in two). *p* values from unpaired, two-tailed Student’s t-test comparing each cell line to parental. (E) Correlation between microtubule growth speed and cytoplasmic EB3 intensity. Simple linear regression is shown (R^2^:0.0065), indicating no correlation. (F) Microtubule growth lifetime and (G) growth length in the indicated cell lines. Data show median ± interquartile range from three independent experiments. *p* values indicate unpaired Mann-Whitney test comparing ranks comparing TTC5 KO to parental cells. (H) Schematic representation of the quantification strategy for α-tubulin and acetylated α-tubulin at the cell cortex. Representative sum-intensity projection images show actin (used to define the cell cortex), α-tubulin, and acetylated α-tubulin channels. *\*\*\*p<*0.001, *\*\*p*<0.01, *\*p<*0.05, *ns* - not significant.

**Extended Data Fig. 2.**
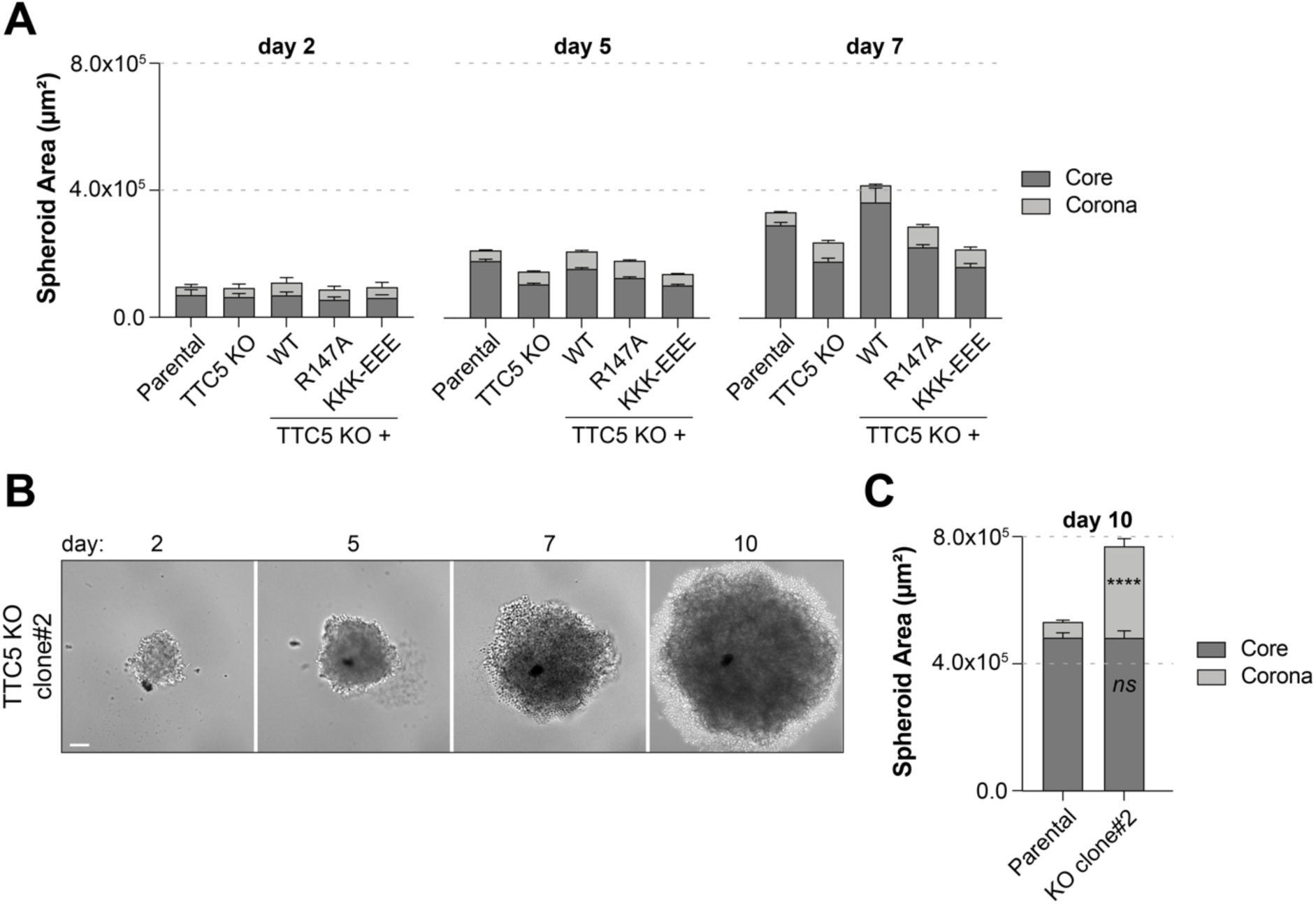
(A) Quantification of spheroid core and corona areas from Fig. 2A at day 2, 5 and 7. (B) Representative brightfield images of 10-day spheroids formed by TTC5 KO (clone#2) cells. Scale bar, 100 μm. (C) Quantification of spheroid core and corona area. Parental spheroids data corresponds to Fig. 2B. Data show mean ± SEM from two independent experiments. *p* values indicate unpaired, two-tailed Student’s t-test comparing TTC5 KO (clone#2) spheroids to parental. *\*\*\*\*p<*0.0001, *ns* - not significant.

**Extended Data Fig. 3.**
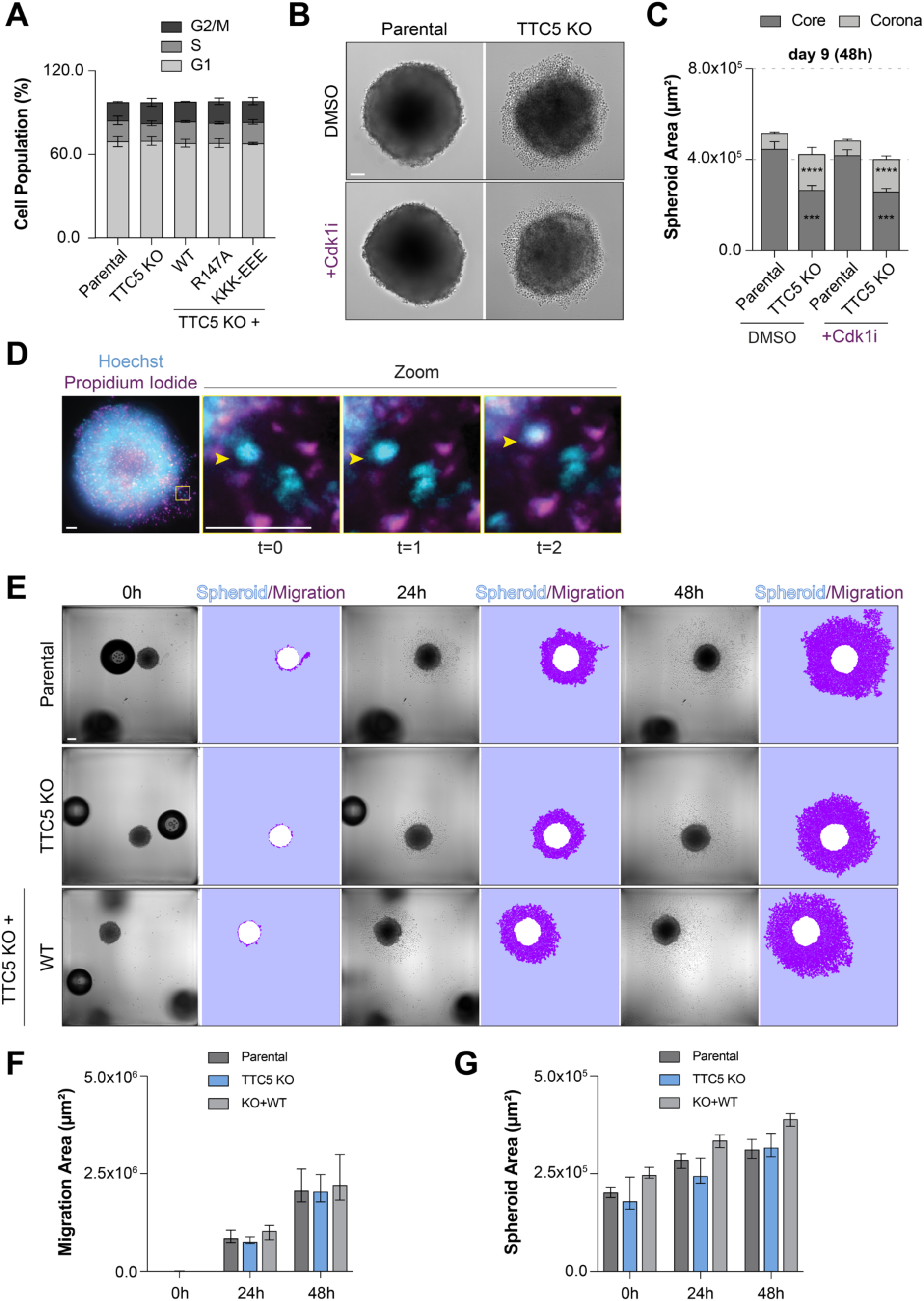
(A) Flow cytometry analysis of cell cycle using PI to assess DNA content. Data represent mean ± SD from two independent experiments. No significant differences were detected using unpaired, two-tailed Student’s t-test between each of the indicated cell lines and parental cell line. (B) Representative images of HeLa parental and TTC5 KO spheroids treated with DMSO or 5 μM Ro-3306 (Cdk1 inhibitor) for 48 hours. (C) Quantification of spheroid core and corona areas at day 9. Data show mean ± SEM from three independent experiments. *p* values from unpaired, two-tailed Student’s t-test comparing TTC5 KO spheroids to parental per condition. (D) Live-cell imaging of TTC5 KO spheroid stained with Hoechst (all nuclei, cyan) and PI (dead nuclei, magenta). Time-lapse every 20 minutes. Yellow arrowhead highlights a cell located at the spheroid corona that dies at t=2 (positive for both Hoechst and PI). Scale bars, 100 μm. (E) Spheroid migration assay performed for 48 hours using HeLa parental, TTC5 KO and TTC5^WT^ lines. Segmentation masks show migration area (magenta) and spheroid area (white). Scale bar, 100 μm. (F) Quantification of migration and (G) spheroid area. Data are mean ± SD from two independent experiments. No significant differences detected using unpaired, two-tailed Student’s t-test for each of the indicated cell lines with the parental cell line as reference. *\*\*\*\*p<*0.0001.

**Extended Data Fig. 4.**
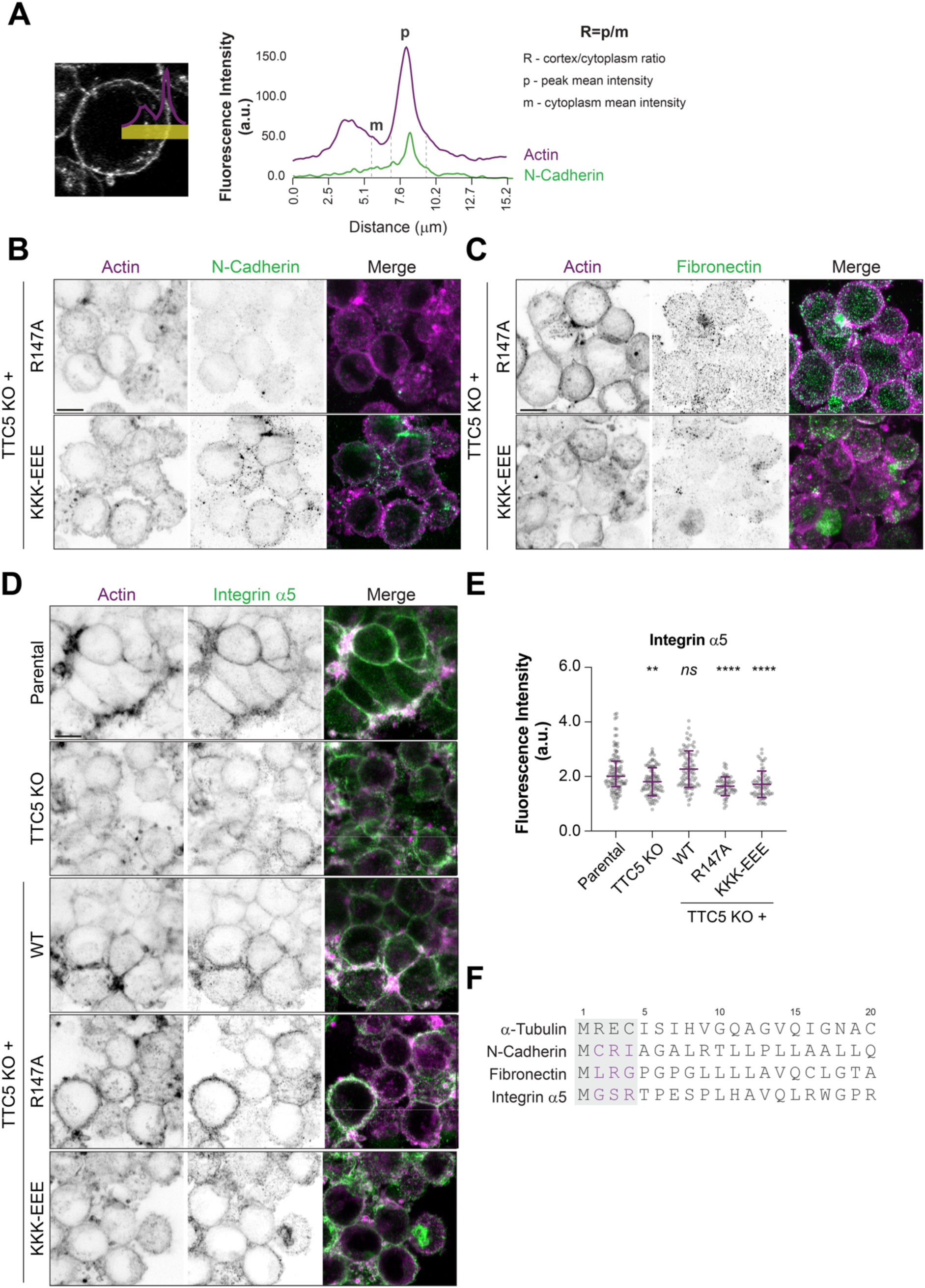
(A) Schematic for calculation of the cortex-to-cytoplasm ratio (R) for actin (magenta) and a protein of interest (e.g., N-Cadherin, green): *p* represents the mean intensity ± 1 μm from cortical peak; *m* represents the mean intensity 1 μm in the cytoplasm, before *p*. (B) Representative confocal images of TTC5^R147A^ and TTC5^KKK-EEE^ spheroids cultured for 10 days counterstained with Phalloidin-647 (magenta, Actin) and immunostained for N-Cadherin (green) or (C) Fibronectin (green). Cortex-to-cytoplasm ratio is shown in Fig. 3B and 3D, respectively. Scale bar, 5 μm. (D) Immunofluorescence of 10-day HeLa parental, TTC5 KO and TTC5^WT^ spheroids stained for Integrin α5 (green) and counterstained with Phalloidin-647 (magenta, Actin). Scale bar, 5 μm. (E) Cortex-to-cytoplasm ratio of Integrin α5 fluorescence intensity. Data show median ± interquartile range from two independent experiments. *p* values indicate unpaired Mann-Whitney test comparing ranks for each of the indicated cell lines with the parental cell line as reference. (F) Sequence alignment of the N-terminal of α-tubulin with those of the adhesion proteins analyzed in this study: N-Cadherin, Fibronectin, and Integrin α5 (1-20 amino acids). The TTC5-recognition motif (MREC), present in α-tubulin, is highlighted in the grey rectangle. This motif is absent in the adhesion proteins (magenta). *\*\*\*\*p<*0.0001, *\*\*p*<0.01, *ns* -not significant.

**Extended Data Fig. 5.**
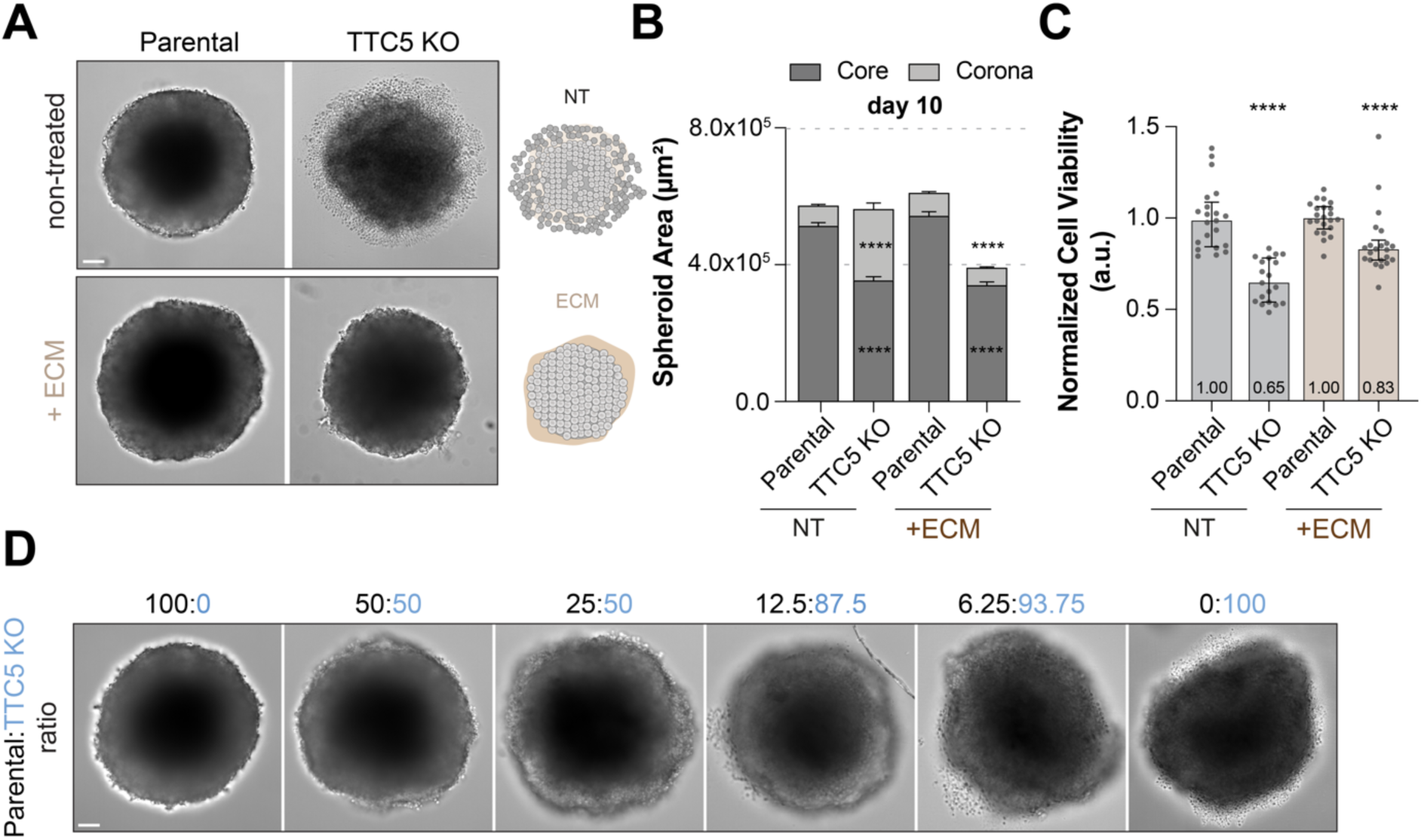
(A) Brightfield images of 10-day HeLa parental and TTC5 KO spheroids grown in normal medium (non-treated, NT) or supplemented with 1 mg/mL Matrigel (+ECM) between days 5 and 7 days. Scale bar, 100 μm. (B) Spheroid core and corona area quantification at day 10, while showing mean ± SEM from three independent experiments. *p* values indicate unpaired, two-tailed Student’s t-test comparing TTC5 KO spheroids to parental per condition. (C) Cell viability determined by ATP content across the indicated cell lines, normalized to parental spheroids per condition. Data show median ± interquartile range from two independent experiments. *p* values indicate unpaired Mann-Whitney test comparing ranks for each group to the respective parental control. (D) Representative images of mixed spheroids formed by parental and TTC5 KO cells at indicated ratios. Spheroid morphology is rescued with as little as 25% of parental cells. *\*\*\*\*p<*0.0001.

**Extended Data Fig. 6.**
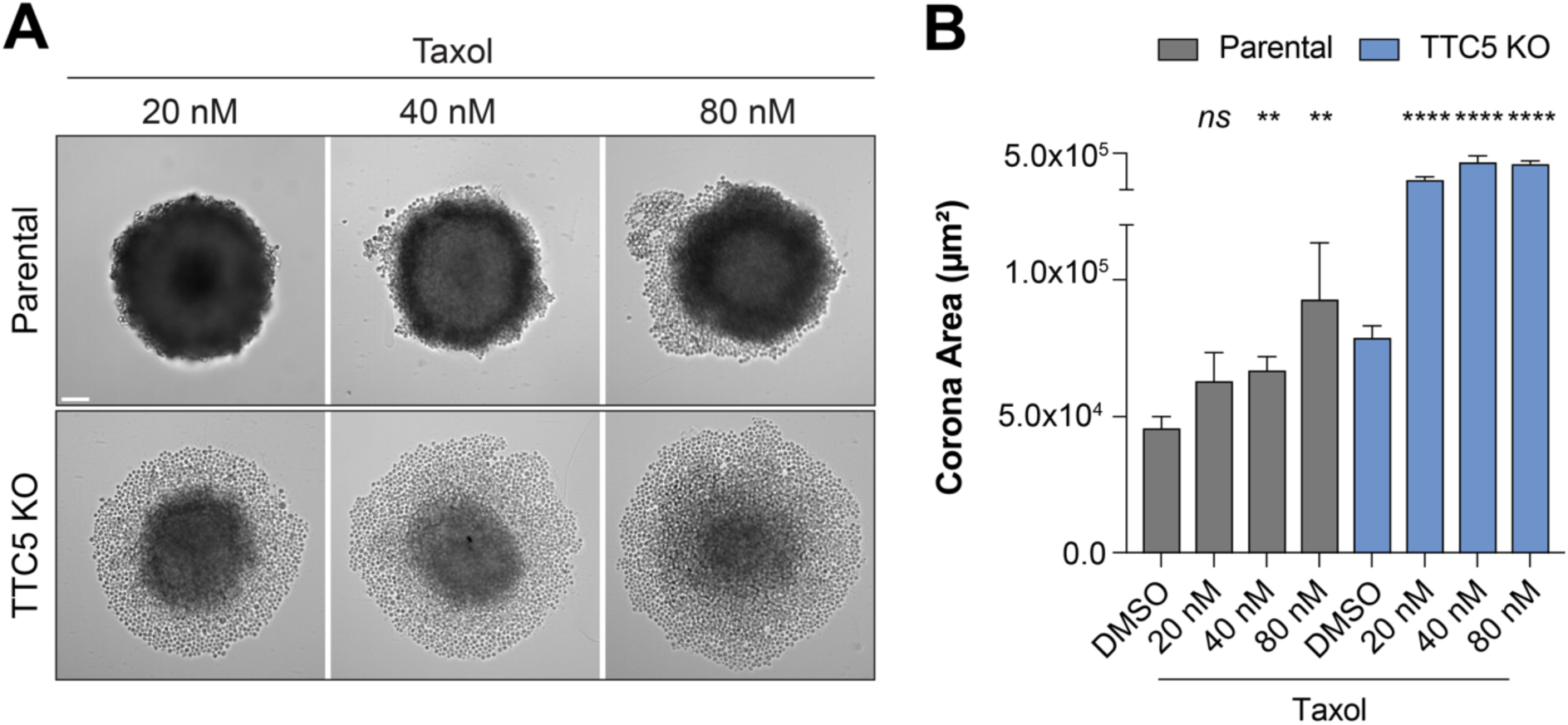
(A) Brightfield images of 7-day spheroids from HeLa parental and TTC5 KO treated with increasing concentrations of taxol (20, 40 and 80 nM) for 48 hours. Corona area expands with increasing taxol doses. Scale bar, 100 μm. (B) Quantification of spheroid areas across the different concentrations at day 7 (48h). DMSO and taxol (40/20 nM, for parental and TTC5 KO, respectively) data are shown in Fig. 4B. Data shown mean ± SEM. *\*\*\*\*p<*0.0001, *\*\*p<*0.01, *ns* - not significant.

